# Single tyrosine mutation in VE-cadherin modulates gene lung expressions: evidence for FOXF1 mediated S1PR1 upregulation to stabilize vessels in mice

**DOI:** 10.1101/2023.07.28.550978

**Authors:** Olivia Garnier, Florian Jeanneret, Aude Durand, Arnold Fertin, Donald Martin, Sarah Berndt, Gilles Carpentier, Christophe Battail, Isabelle Vilgrain

## Abstract

**Rationale:** Phosphorylation-dephosphorylation are processes involved in the adhesion of endothelial cells (ECs) to maintain vascular integrity in adults. VE-cadherin is a target for Src-mediated Y^685^ phosphorylation, identified in highly vascularized human glioblastoma where it is involved in the abnormal feature of tumor blood vessels.

**Objective:** We aimed at understanding the molecular mechanisms through which Y^685^F-VE-cadherin triggers S1PR1 gene expression and stabilizes lung vessels in adult mice.

**Methods and Results:** We compared lung ECs from a knock-in (KI) mouse carrying a point mutation in VE-cadherin (Tyr 685 to Phe) to Wild type. Analysis of EC parameters showed a difference in the migratory rate was between ECs from KI (22.45% ± 5.207) and WT (13.24% ± 5.17) (p-value=0.034). The direct adhesion of ECs from KI mice to fibronectin was significantly higher (37.625 ± 9.23) than that of the WT (26.8 ± 3.258, p-value=0.012). In the fibrin bead assay, ECs from KI showed a weaker angiogenic response. The transcriptome of mutated ECs showed that 884 genes were dysregulated of which 766 genes were downregulated and 118 genes were upregulated. The Gene Ontology Enrichment showed that most of the genes were related to cell-cell adhesion and angiogenesis. Focusing on angiogenic genes, we found that Sphingosine-1-phosphate-receptor was a gene upregulated in mutated ECs which was confirmed by RT-PCR and westernblotting. Mechanistically, chromatin immunoprecipitation assay (CHIPS) demonstrated that FOXF1 directly bound to the S1pr1 promoter 7 fold greater than WT. As a consequence, VE-cadherin at the membrane was higher in the mutant vs WT (100 ± 6.52 for WT vs 189.7 ± 21.06 for KI (p-value 0.0001). Finally, lung morphometric analysis showed less vessels and vascular remodeling with no fibrosis in mutated mice.

**Conclusions:** These data extend our knowledge on pY-VE-cadherin mediated pathological angiogenesis and provide new therapeutic opportunities to vascular normalization through pharmacological inhibition of the Y685-VE-cadherin phosphorylation.

## Introduction

Endothelial cells (ECs) are lining the lumen of blood vessels and orchestrate important processes such as vasomotor tone, coagulation, permeability, tissue vascularization and immune response (1). The vascular network responds to the subsequent metabolic changes in a rhythmic pattern with the change in the physiology of several organs specially those which are subjected to hormonal control (2). While ECs remain quiescent throughout adulthood, the activation of endothelium occurs during patho/physiological conditions and lead to either endothelial activation or to angiogenesis (3). The process of angiogenesis is dynamically adjusted with several other growth factors and controlled by suitable regulators, such as vascular endothelial growth factor (VEGF) (4). Endothelial adherens junctions are adhesive intercellular contacts that are crucial for the maintenance and regulation of normal vascular function. Alterations in adherens junction assembly influence endothelial cell motility, vascular morphogenesis and permeability. The inability of ECs to fulfill their physiological functions is a key feature of multiple prevalent diseases including hypertension, atherosclerosis, diabetes and cancer. More precisely, we reported that endothelial cell dysfunction occurs in several cancers including renal cell carcinoma, glioblastoma and breast cancer (5–7).

VE-cadherin is an adhesive protein exclusively expressed in ECs whose extracellular domain has an important role in endothelial cell-cell adhesion, whereas the cytoplasmic tail interacts with the actin cytoskeleton via three proteins of the armadillo family, called β-catenin, plakoglobin, and p120 to ensure junctional strength (8). The cytoplasmic domain of VE-cadherin was shown to regulate endothelial protrusive activity in vitro, suggesting that this domain may play a prominent part in invasive processes (9). Tyrosine phosphorylation of the cadherin–catenin complex has been proposed as a mechanism that regulates the stability of cell–cell junctions (10). Interestingly, the cytoplasmic domain of VE-cadherin contains nine tyrosine that represent potential target sites for tyrosine kinases. VE-cadherin tyrosine phosphorylation has been reported in ECs under several conditions including stimulation by VEGF. The non-receptor tyrosine kinase of the Src family have an important role in these processes and Src was shown to be associated with VE-cadherin in vivo (11)) and to induce phosphorylation at site Y685, which correspond to the consensus site (LY^685^ AQV) which fits the YxxV/I/L motif, known for Src kinases (12,13). In attempt to determine whether human angiogenic tumor tissues have some phosphorylated VE-cadherin molecules in the capillary network, we have investigated glioblastoma because it is a highly vascularized tumor (14, 6). We found that the phosphorylation of the site of Y^685^ in VE-cadherin from human surgical tissues. Due to the importance of angiogenesis in cancer, it was crucial to determine the role of this tyrosine in VE-cadherin. For that purpose, we developed a knock-in mouse (KI) carrying a tyrosine-to phenylalanine point mutation of VE-cadherin Y^685^ (VE-Y^685^F) to study the phenotype of the homozygous mice (15,16). The single point mutation on one amino acid is not lethal but the KI mice rather developed oedema in organs subjected to hormonal regulation such as ovary and uterus (16). These results as well as other group datas suggested that the Y^685^ was responsible of increased permeability (17). Our ultrastructural analysis of endothelial junctions of Y^685^F-VE-cadherin ovary showed a slight increase in the gap between two adjacent ECs, suggesting that mutation Y^685^F may cause nanomodifications in the structure of endothelial junctions that may destabilize them (15). Because VE-cadherin is also linked to the actin cytoskeleton, the impact of the mutation might affect the binding of its intracytoplasmic partners including β-catenin which role in cell-cell adhesion control has been well documented (18). In addition, β-catenin can also translocate to the nucleus and acts a as a transcriptional activator (19). Given that VE-cadherin is exclusively expressed in ECs, it was reasonable to hypothesize that a single mutation in the protein would affect the global profiling of transcript (transcriptomics) and could open new directions to unravel the molecular mechanisms that underlie EC functions. In the present report, we compared lung ECs from KI to wild type ECs. We demonstrate that their angiogenic properties were modified. In addition, we demonstrated mechanistically that β-catenin is dissociated from its point of anchorage in VE-cadherin since the mutant is phosphorylated on Y^731^. Therefore, β-catenin translocates to the nucleus to activate S1PR1, which is a direct transcriptional target of the FOXF1 in lung ECs from KI mice compared to WT. As a consequence, S1PR1 maintained VE-cadherin at the membrane of ECs and protected the lung against pulmonary fibrosis. Altogether, our findings suggest that the effect of a single mutation of VE-cadherin participates at least to pulmonary vascular remodeling and thus this phosphorylation events on Y^685^ of VE-cadherin could lead to fibrosis-associated EC with high permeability which contributes to lung inflammation.

## METHODS

### Animals

Four-week-old male wild-type (WT) C57BL/6 mice and four-week-old male transgenic Y685F-VE-cadherin (KI) C57BL/6 mice were used for the experiments and were generated as previously described (15). Mice were maintained in a conventional animal facility, on a 12-hour light/12-hour dark cycle. Food and water were available ad libitum. All procedures were carried out in compliance with the principles and guidelines established by the National Institute of Medical Research (INSERM) and approved by the Institutional by the Institutional Ethical Committee for animal experimentation (Comité d’Ethique). Animals were anesthetized by an intraperitoneal injection of ketamine (100 mg/kg) and xylazine (10 mg/kg).

### Reagents

Reagents were purchased from several source: collagenase type I (10114532, Fisher Scientific), DMEM (4.5g/L Glucose) (11965092, Gibco), EGM-2 BulletKit (CC-3156 & CC-4176, Lonza). Pierce BCA dosage Assay kit (23225, ThermoFisher) was used for protein concentration of lung and isolated ECs extracts. Protein G Sepharose 4B fast flow (P3296, Sigma Aldrich) was used for the Co-immunoprecipitation and ChIP assay. For RNA extraction Nucleospin® RNA (740955.50, Machery-Nagel) was used. For cDNA synthesis, 5x iScript Reaction Mix (1708890, Biorad) was used. For qPCR, SsoAdvanced Universal SYBR Green Supermix (1725270, Biorad) was purchased. For PCR, EconoTaq Plus Greeen (30033, Lucigen) was purchased. Mouse Angiogenesis array kit (ARY015, R&D system) was purchased for protein array. For Masson trichrome staining, a special kit (H15) and hematoxylin Gill (GHS332) and eosin (HT110232) were purchased at Sigma Aldrich.

### Antibodies

Commercially available antibodies were purchased from several sources: Antibodies used for the FACS: CD-31 (PE) antibodies were from Becton-Dickinson, CD45R (53-0452-82, Fisher scientific). For Miltenyi cell isolation: CD45 microbeads (130-052-301, Miltenyi), CD31 microbeads (130-097-418, Miltenyi) and MS Separate column (130-042-201, Miltenyi) were used. Anti-VE-cadherin antibody (AF1002) and anti FOXF1 antibody (AF4798) were purchased from R&D system). Anti S1PR1 antibody was from (SAB4500687, Sigma-Aldrich). Anti-VEGFR2 antibody (sc-6251), anti-VEGF antibody (sc-7269), anti-Src antibody (sc-18), anti-VE-cadherin antibody (sc-19) were from Santa-Cruz. Anti-pY^1175^-VEGFR2 antibody (D5B11), and anti Ill-catenin antibody (9562) were purchased from Cell signaling. Anti-pY^418^-Src (LF-PA20465) was from Ab Frontier. Anti-pY^658^-VE-cadherin antibody (44-1144G), and anti-pY^731^-VE-cadherin antibody (441145G) were from Invitrogen. Cy3 Donkey anti-Goat (705-166-147) secondary antibody was from Jackson.

### Isolation of ECs using fluorescent activated cell sorter (FACS)

4 week-old male mice were sacrificed and the lungs were immediately placed in cold sterile PBS (Ca-/Mg-). The tissues were shredded using scalpel into small pieces then transferred into a previously autoclaved Erlenmeyer flask containing collagenase type 1. Collagenase type I was prepared in DMEM (4.5g/L Glucose) with no serum at a concentration of 3mg/mL and filtered with a 0.22µm filter before the start of the experiment. The digestion last 45 minutes at 37°C and every 15 minutes, the Erlenmeyer was shaken to improve the collagenase type I activity. The digestion was stopped by adding an equal volume of DMEM (4.5 g/L Glucose) with 20% FBS. The cells suspension was passed through a 10 mL syringe with a 18G needle, then filtrated with the following filters: 100, 70µm and 40 µm diameter to obtain a homogenous cell suspension. The cells were then centrifuged 5 minutes at 1200 rpm and resuspended in DMEM (4.5 g/L Glucose) with 20% FBS for cell counting. The solution was centrifuged again 5 min at 1200 rpm to resuspend the cell pellet at 10.10^6^ cells/mL in PBS(Ca+/Mg+) / BSA 2%. The cell suspension was distributed into separate falcon tubes for immunostaining: one as a negative control, one with only CD45-Alexa 488 antibody, one with the antibody CD31-PE only. The samples of interest was stained with the antibody CD31-PE and CD45-Alexa 488. The tubes were incubated for 1 hour in the dark at 4°C and shaken every 10 minutes. At the end of the incubation, the cells were washed with 10 mL of PBS(Ca+/Mg+) / BSA 2% and centrifuged 5 min at 1200 rpm. The supernatant was removed carefully and 500µl of DMEM (4.5 g/L Glucose) without serum was added in each tube. The cell solution was filtered through FACS tube and the labeled CD31 positive and CD45 negative were collected in DMEM (4.5 g/L Glucose) with 20% FBS. The stained cells were washed twice, resuspended in PBS and analysed in a FACScaliber flow cytometer (Becton Dickinson).

### Isolation of the ECs from the lung with Miltenyi method

4 week-old male mice were sacrificed. Mouse lungs were harvested and placed in cold PBS (Ca-/Mg-) to be cut in small pieces and digested with 4 mL of collagenase I (3mg/mL) in DMEM (4.5g/L Glucose) with no serum at 37°C for 45 min. The digestion was stopped by adding 20% FBS. The cells suspension was passed through a 10 mL syringe with a 18G needle followed by several filters: 100, 70µm and 40 µm diameter. The cell suspension was centrifuged 5 minutes at 1200 rpm and resuspended in 4 mL of complete medium before another centrifugation for 5 min at 1200 rpm. The cell pellet was resuspended in PBS (Ca-/Mg-) /FBS 2% (90µL of PBS-/-/FBS 2% up to 10^7^ cells) and CD45 mouse microbeads were added according to the manufacturer instructions (10µl up to 10^7^ cells) and incubated for 15 min at 4°C. After a wash with PBS (Ca-/Mg-),FBS 2%, the cells were centrifuged 5 min at 1200 rpm and resuspended in 500µL of PBS (Ca-/Mg-), /FBS 2%, and pass through MS column. After 3 washes with PBS(Ca-/Mg-) /FBS 2%, the CD45-fraction was kept, centrifuged and resuspended with the CD31 Mouse Microbeads for 15 minutes at 4°C in the dark. After washing, centrifugation 5 min at 1200 rpm, the cells were passed through MS column as described before. The CD 31 positive fraction was collected by adding 1 mL of PBS (Ca-/Mg-) / FBS 2% into the column and flushed firmly. The cell suspension was seeded with EGM-2 onto fibronectin-coated wells (300 000 cells per well). The cells were cultured 5 days before experiments or used for RNA isolation.

### Quantitative real time RT-PCR (qRT-PCR)

#### mRNA isolation from the endothelial cells

Nucleospin® RNA kit was used for the isolation of RNA from the cultured cells. The protocol was followed exactly according to the manufacturer. RNA was eluted with 40µL ofautoclaved distilled water The purity of RNA was checked and quantified with a NanoDrop spectrophotometer (Thermo Fisher, Massachusetts, USA)

#### Reverse transcription of RNA to cDNA

The conversion of RNA isolated from the ECs to cDNA was done using iScript cDNA Synthesis Kit. The sample reaction was prepared following the manufacturer instructions. The complete reaction mix was incubated in a thermal cycler in the following protocol: 5 minutes at 25°C, 20 minutes at 46°C and 1 minute at 95°C.

#### Quantitative Polymerase Chain Reaction (qPCR)

SsoAdvanced Universal SYBR Green Supermix was used for cDNA amplification with different genes primers (see table S1) on the CFX96 Real-Time PCR Detection System using the following conditions: 95°C 30 seconds/ 95°C 5 seconds; 60 °C 15 seconds for 39 cycles, 95°C for 10 seconds, 65°C for 5 seconds and 65°c for 5 seconds.

#### Polymerase Chain Reaction PCR

DNA samples from ChIP were analyzed by qPCR. DNA were amplified with FOXF1 binding promoter primers (see table S1) and EconoTaq Plus Greeen according the following cycles: 3 minutes at 95°C, 34 cycles: 95°C 15 seconds; 62°C 15 seconds; 72°C 20 seconds, 72°C 10 minutes. The samples were loaded onto the agarose gel for analysis.

The relative level of RNA was normalized to housekeeping gene *hprt1*. Fold change was calculated using the 2((−ΔΔCt) method) (20).

#### Transcriptomic analysis

The ECs from the lung was isolated with the FACS method as described before and the cell pellet were frozen in nitrogen liquid and sent to GENEWIZ for RNA-seq (Genomics European Headquarters Bahnhofstr. 8 04158 Leipzig Germany).

### RNA Extraction, Library Preparation, NovaSeq Sequencing, and Standard RNA-Seq analysis

Total RNA was extracted using Qiagen RNeasy Mini kit following manufacturer’s instructions (Qiagen, Hilden, Germany). RNA samples were quantified using Qubit 4.0 Fluorometer (Life Technologies, Carlsbad, CA, USA) and RNA integrity was checked with RNA Kit on Agilent 5300 Fragment Analyzer (Agilent Technologies, Palo Alto, CA, USA).

RNA sequencing libraries were prepared using the NEBNext Ultra RNA Library Prep Kit for Illumina following manufacturer’s instructions (NEB, Ipswich, MA, USA). Briefly, mRNAs were first enriched with Oligo(dT) beads. Enriched mRNAs were fragmented for 15 minutes at 94 °C. First strand and second strand cDNAs were subsequently synthesized. cDNA fragments were end repaired and adenylated at 3’ends, and universal adapters were ligated to cDNA fragments, followed by index addition and library enrichment by limited-cycle PCR. Sequencing libraries were validated using NGS Kit on the Agilent 5300 Fragment Analyzer (Agilent Technologies, Palo Alto, CA, USA), and quantified by using Qubit 4.0 Fluorometer (Invitrogen, Carlsbad, CA).

The sequencing libraries were multiplexed and loaded on the flowcell on the Illumina NovaSeq 6000 instrument according to manufacturer’s instructions. The samples were sequenced using a 2×150 Pair-End (PE) configuration v1.5. Image analysis and base calling were conducted by the NovaSeq Control Software (NCS). Raw sequence data (.bcl files) generated from Illumina NovaSeq was converted into fastq files and de-multiplexed using Illumina bcl2fastq program version 2.20. One mismatch was allowed for index sequence identification.

After investigating the quality of the raw data, sequence reads were trimmed to remove possible adapter sequences and nucleotides with poor quality using Trimmomatic v.0.36. The trimmed reads were mapped to the mouse reference genome available on ENSEMBL using the STAR aligner v.2.5.2b. BAM files were generated as a result of this step. Unique gene hit counts were calculated by using feature Counts from the Subread package v.1.5.2. Only unique reads that fell within exon regions were counted.

After extraction of gene hit counts, the gene hit counts table was used for downstream differential expression analysis. Using DESeq2, a comparison of gene expression between the groups of samples was performed. The Wald test was used to generate P values and Log2 fold changes. Genes with adjusted P values < 0.05 and absolute log2 fold changes >1 were called as differentially expressed genes for each comparison. Gene ontology analysis was performed on the statistically significant set of genes by implementing the software GeneSCF. The goa_Mus musculus GO list was used to cluster the set of genes based on their biological process and determine their statistical significance.

A PCA analysis was performed using the “plotPCA” function within the DESeq2 R package. The plot shows the samples in a 2D plane spanned by their first two principal components. The top 500 genes, selected by highest row variance, were used to generate the plot.

The functional enrichment analysis was performed based on the “Biological Process” ontologies from the Gene Ontology database (GO.db R package version 3.16.0). ORA and GSEA were computed using the Bioconductor/R package clusterProfiler (version 4.6.0). Gene sets with a corrected p-value (Benjamini-Hochberg correction method) < 0.05 were considered significant. Protein-protein networks were built using the STRING-DB web software (https://string-db.org/) using significant DEGs found in cell-cell adhesion or angiogenesis ontologies. Statistical analysis were performed using R (version 4.2.2).

### Cell migration assay

The freshly isolated ECs were immediately seeded at 200 000 cells / well in 96 well plates pre-coated with fibronectin and grown to confluency in the EGM-2 medium. The day of the wound assay, the cells were washed with PBS (Ca+/Mg+) and the scratch was performed with the Incucyte® Cell Migration Kit (Sartorius). The cells were washed carefully twice with PBS (Ca+/Mg+) then DMEM (4.5G/L Glucose) containing 1.5% FBS was added. The full time course migration was performed for 20h in the Incucyte Zoom live-cell analysis system (Sartorius) and the analysis of the gap area of the wound was performed by the Incucyte Basic Analyzer software.

### 3D fibrin beads gel assay in vitro

3D fibrin gel bead assay was performed as described previously (21). Isolated ECs were mixed with Cytodex 3 microcarrier beads at the concentration of 800 ECs per beads. In 1 mL of warm EGM-2 medium. Beads and ECs were then co-incubated in a humidified incubator at 37 °C and 5% CO2 and gently manually shaken every 20 min for 4 h to allow cell adherence to the bead surface. After 4 hours, they were transferred into 25cm^2^ flask in 5 mL of EGM-2 medium overnight. The following day, the coated beads were resuspended in 2.5mg/ml fibrinogen solution, 1µg/mL aprotinin and 10ng/mL VEGF at concentration of ̴ 500 beads/ml. 12µL of thrombin 10U/mL and 288µL of the mix fibrinogen/beads suspension per well of 24-well plate. The plate was incubated 15 minutes in a humidified incubator at 37 °C and 5% CO2 until clot formation. 500µL of EGM-2 supplemented with angiopoetin 1 (250µg/mL), hGF (100ng/mL), IGFBP3(50ng/mL), VEGF(20ng/mL), PDGF(10ng/mL), EGF(100ng/mL) (21). After 4 days of incubation, the angiogenic sprouting were imaged with the Axioobserver Z1 Zeiss software at the magnification of X5. Quantification of morphometrical parameters of the capillary network was performed using a program developed for the Image J software (22). This plugin is an extension of the “Angiogenesis Analyzer” for Image J written in the macro language of Image J. Representative parameters measured were number of anchorage per junctions, number of junctions, total master segments and number of meshes per beads.

### Adhesion assay

Isolated ECs were seeded in 96 well-plates pre-coated with fibronectin in EGM-2 (1500 cells/well). After 30 min of incubation, the cells were washed with PBS (Ca+/Mg+), fixed with PFA 4% and permeabilized with 0.2% Triton X100 for 20 minutes at room temperature. The nuclei were stained with Hoescht (1/400) for 5 minutes in PBS. After 3 washes with PBS (Ca+/Mg+), the cells were imaged with the Axio-observer Z1 Zeiss software at the magnification of X5. The number of nuclei were counted with Image-J software (NIH, Bethesda, MD).

### Proliferation assay

Isolated ECs were seeded onto 96 well-plates pre-coated with fibronectin in EGM-2 and the proliferation assay was performed for 60 hours in the Incucyte proliferation assay according to (21) (Sartorius). The analysis of the cell proliferation was performed by the Incucyte Basic Analyzer software.

### Immunofluorescence

Isolated ECs were cultured 4-7 days with EBM-2 and then washed with PBS (Ca+/Mg+). The cells were fixed with PFA 4% and permeabilized Triton 0.2% 20 minutes at room temperature. The cells were washed twice for 5 minutes with PBS (Ca+/Mg+) and blocked with PBS (Ca+/Mg+) / BSA 3% for 30 minutes at room temperature. The cells were incubated with the mouse VE-cadherin primary antibody in PBS (Ca+/Mg+) / BSA 3% for 1h at room temperature with gentle agitation. After two washes with PBS (Ca+/Mg+), Tween 0.1% for 5 minutes, Cy3 Donkey anti-Goat antibody (1/200) was added in the dark 1h at room temperature under gentle agitation. The nuclei were stained with Hoechst in PBS for 5 minutes at room temperature in the dark before the slides were mounted. The slides were imaged with the Axio Imager M2 Zeiss microscope (magnification X63). VE-cadherin staining was quantified by measuring the staining area at the membrane and the integrated density of the staining using Image-J software macro (NIH, Bethesda, MD). Scale bars are 20 μm.

### Chromatin immunoprecipitation assay (ChIP)

The ChIP protocol was kindly given by Dr Marco Saponaro. Briefly, 4 week-old male mice WT and KI was sacrificed and the lungs were harvested and placed immediately in cold PBS (Ca-/Mg-). Immersed tissues were shredded using scalpel into small pieces and digested with 4 mL of collagenase I (3mg/mL) in DMEM (4.5g/L Glucose) with no serum at 37°C for 45 min. Every 15 minutes, the tube was shaken to improve the collagenase I activity. The digestion is stopped with 20% FBS. The cells suspension was pass through a 10 mL syringe with a 18G needle and through several filters: 100, 70µm filters followed by centrifugation at 1200 rpm for 5 minutes. The supernatant was carefully removed and resuspended in PBS. Formaldehyde was added to 1% final concentration and incubated in rotation at room temperature for 15 minutes. Glycine (125mM) was immediately added and followed by 5 minutes incubation at room temperature under agitation. The solution was centrifuged 2 minutes at 14000 rpm and washed twice with ice-cold PBS for 2 minutes at 14000 rpm. Cells were lysed with ChIP buffer (5mM HEPES, pH 8.0, 85mM KCL, 0.5% NP-40 and leupeptin (10µg/mL), Na3VO4 (2mM)), and incubated 5 minutes on ice. The cell solution was centrifuged 5 minutes at 3900 x g at 4°C to obtain the nuclei pellet. The nuclei were lysed with ChIP nuclear lysis buffer (50mM Tris/HCL pH 8.1, 10mM EDTA, 1% SDS + leupeptin (10µg/mL) and Na3VO4 (2mM)) and incubated 5 minutes on ice. Nuclear lysate was sheared by using an ice-water bath embedded sonication system at high power, 30 seconds on, 30 seconds off mode for 10 minutes. Sonicated chromatin was cleared by centrifugation at 20,000g for 10 min at 4°C. Chromatin was diluted 1:5 with ChIP dilution buffer (0.01% SDS, 1.1% Triton X-100, 1.2mM EDTA (pH 8.0), 16.7mM Tris/HCL pH 8.1, 167mM NaCL, leupeptin (10µg/mL), Na3VO4 (2mM)). The sample was cleared with 30µL of Sepharose beads G for 30 minutes under agitation at 4°C and centrifugation at 6000 rpm 2 minutes at 4°C. The supernatant was transferred into a new tube and 2.5µg of FOXF1 antibody was added overnight at 4°C under agitation. Protein G Sepharose beads were added into the tubes and incubated 3h at 4°C under agitation. Washes are performed sequentially, doing them two times with 1ml of each of the following buffers: ChIP low salt buffer(0.1% SDS, 1% Triton X-100, 2mM EDTA, 20mM Tris/HCL pH 8.1, 150mM NaCl) and ChIP high salt buffer (0.1% SDS, 1% Triton X-100, 2mM EDTA, 20mM Tris-HCL pH 8.1, 500mM NaCL) followed by ChIP LiCl buffer (10mM Tris-HCl pH 8.0, 250mM LiCl, 1%NP40, 1% deoxycholic acid, 1mM EDTA). A final wash was made with TE buffer (pH=8) and centrifuged at 3000 rpm for 3 minutes. The pellet was resuspended in 40μL elution buffer (50mM Tris/HCL pH 8.0, 10mM EDTA, 1%SDS) with RNAse 10mg/L and incubated overnight at 65°C. The sample was centrifuged 1 minute at 12000rpm and the supernatant was transferred into a fresh tube where 120µL TE/1%SDS was added with proteinase K (20mg/mL) and glycogen (10mg/mL) whereas the pellet were resuspended in Laemmli 2X for western blot analysis. The chromatin solution was incubated for 2 hours at 37°C. An exact equal volume of Phenol/Chloroform/Isoamilalchol 25:24:1 pH 8.0 was added and mixed by inverting the tube until the phases are completely mixed. Then, the tube was centrifuged at 12000 rpm 5 minutes and the upper layer was collected into a new tube where ethanol 100% and 5M NaCL were added. The sample was stored at −20°C for 1 hour. It was centrifuged again 10 minutes at 12000 rpm. The pellet was resuspended in ethanol 70% and centrifuged 10 minutes 1200 rpm. The supernatant was retrieved as the pellet let completely dry before adding TE buffer to perform the PCR.

### Histological sectioning of lungs and staining

For all histological procedures, the lungs were inflated to full capacity and fixed by intratracheal instillation of formalin 10% (1 mL), immersed in formalin overnight, and then embedded in paraffin. Briefly, mice were sacrificed and lung were harvested quickly and transferred in ethanol 90% 2 hours at room temperature, then in ethanol 100% 3 hours still at room temperature. After 4 times in methylcyclohexane for 45 minutes at room temperature, the lungs were placed in paraffin/methylcyclohexane (1:1) at 60°C for 24 hours. Lungs were transferred into fresh paraffin every day for 4 days, and let to dry at room temperature. Paraffin embedded lungs were sagittally sectioned at 5 μm and kept at 60°C for an hour. Sections were stained with Harris’s hematoxylin/eosin or Masson’s trichrome stain for general morphology and morphometric analysis. For Hematoxylin and eosin: the sections were washed with xylene before serially washing with 100% ethanol for 5 minutes 3 times. The sections were incubated into the stain for 2 minutes, then rinsed with water and treated with eosin directly for 3 minutes, washed with ethanol then xylene before mounting with Aquamount.

The trichrome staining was performed according to the manufacturer’s instructions. Briefly, the sections were treated with warm (56°C) Bouin’s solution for 15 min and stained with Weigert’s hematoxylin for 2 min, then with Biebrich Fuchsin for 5 min, rinsed, and immersed in phosphomolybdic acid for 5 min and stained with Aniline blue for 5 min, and rinsed in distilled water for 2 min, immersed in 1% acetic acid for 2 minutes. Finally, the sections were dehydrated in ethanol, left to air dry, and mounted for Axioplan microscrope. Masson’s trichrome staining showed collagen fibers in blue. Image post analysis was performed with Image-J software (NIH, Bethesda, MD).

### Quantification of immunochemistry

Pulmonary arteries from 4 week-old mice (4 WT and 4 KI) were analyzed. Measurements of the vessel number were taken for each slide corresponding to each mouse lung. Only arteries between 50-150µm of luminal diameter were selected for analysis. We used Image J for measurement of total vessel area (µm^2^), luminal area (µm^2^), and inner circumference (µm). Medial area (µm2) was calculated as the difference between total and luminal areas. Standard medial thickness (µm) was calculated as the ratio of medial area to inner circumference (23).

### Analysis of VE-cadherin Immunostaining

VE-cadherin signal and nuclei were segmented by first denoising images with a Gaussian filter (σ = 1), and then denoised images were thresholded with the triangle algorithm. The mean of VE-cadherin signal was computed over its segmentation mask, and the total area of nuclei was computed from Hoescht channel mask. In order to obtain an invariant measure of VE-cadherin signal regarding to both cell number and cell size, the mean VE-cadherin was divided by the total area of nuclei (24);

### Western blot analysis and immunoprecipitation

Protein extracts from lung and isolated ECs were analyzed by Western blot analysis as previously described^8^. The antibobies used in the present study were: VE-cadherin (AF1002) (1/500), pY^731^VE-cadherin (1/500), pY^658^VE-cadherin (1/500), pY^416^Src (1/1000), Src (1/500), β-catenin (1/1000), VEGFR2 (1/500), pY^1173^VEGFR2 (1/500), VEGFA (1/500), S1PR1 (1/250). Protein expression was normalized with actin or vinculin expression level as indicated. The results are presented as mean ± SEM.

For immunoprecipitation, 100μg of proteins were incubated with 1µg of VE-cadherin antibody (sc-19) overnight. Then the samples were incubated with protein G Sepharose for 3 hours, then washed run onto SDS-PAGE 10%. The proteins were then transferred onto a 0.45μm nitrocellulose membrane. Membranes were blocked with milk, incubated with primary antibodies and corresponding HRP-conjugated secondary antibodies. Immunoreactive proteins were visualized by Bio-Rad ChemiDoc™ with ECL reagent (Bio-Rad Laboratories) and were quantified using densitometry with Image-J software (NIH, Bethesda, MD).

### Angiogenesis protein array

Analyzis of the protein expression in the broncho-alveolar lavage of mice was performed with the Mouse Angiogenesis Array Kit. Briefly, the BAL from 4 week-old male mice of each genotype was centrifuged at 1000g and the supernatant was concentrated on Amicon Ultra-2 - Ultracel-10 membrane, 10 kDa and protein concentrations was determined using the Micro BCA Protein Assay Kit. The same amount of protein concentration for each mouse genotype was analyzed using the Angiogenesis assay according the manufacturer instructions (4 membranes). Immunoreactive proteins were visualized by Bio-Rad ChemiDoc™ with ECL reagent (Bio-Rad Laboratories) and were quantified using densitometry with Image-J software (NIH, Bethesda, MD).

### Statistical analysis

At least three to five mice per group were used in each set of experiments. The experiments were performed at least three times under identical conditions with comparable results. Data are represented as individual values plus plus means ± SD. The sample size, what each “n” represents, the statistical tests used and the result of the statistical test are indicated in each respective figure legend. Statistical analyses were performed using Prism v8.0 (GraphPad) software. Differences between two different groups were analyzed by unpaired Student’s t-test. A probability value <0.05 indicated by an asterisk* showed a significant difference between 2 groups. Non significant difference was indicated by ns.

For immunostaining and immunofluorescence images, assessment of the differences were performed using boxplot and scatter plot in R along with Kruskal-Wallis rank sum test and chi-square test to determine the p-value.

## RESULTS

### Isolation of lung ECs for transcriptomic analysis

We studied ECs from mice lung to avoid the considerable structural, phenotypic, and functional heterogeneity of endothelium depending on the tissue (1). The mice used in this study were referred as to wild type (WT) and the knock-in mouse carrying a tyrosine-to phenylalanine point mutation of VE-cadherin Y685 (KI). Because in all the body, each vascular bed has unique structural and functional properties, we chosen pulmonary ECs from 4 week-old male mice as the lung is highly vascularized (11). Furthermore, we previously showed that although VE-cadherin could be detected in all mouse tissues, there was a dramatic expression in lung (11). Thus, the ECs were isolated from the lungs of (Fig 1A). We performed flow cytometry analyses of cells from both genotypes mice after collagenase dissociation and single-cell suspensions staining for surface antigens (CD31 and CD 45 labelling). CD31 is a uniform marker of endothelial cells, resistant to collagenase treatment (25). The percentage of ECs was obtained using light scatter and fluorescence characteristics. The antibody to CD31 recognised a distinct population (Fig. 1A), whereas cells incubated with non immune rat immunoglobulins harboured background staining (Neg). The percentages of CD31-positive cells in lung tissues were highly reproducible; variations were found to be less than 5% of the mean when three independent measurements were performed on one lung. Representative examples of flow cytometry analyses of cells from lungs are shown in Fig. 1A. Mean percentages of CD31-positive cells were remarkably stable (*n* = 4 independent lungs in each group). We next wanted to determine the expression of cell surface endothelial markers. Total RNA was reverse-transcribed and amplified using exponential non saturating conditions with specific oligonucleotides for Cdh5, and CD31. PCR products were separated on ethidium bromide-stained 2.5% agarose gel. α-SMA, and ptprc were used to determine the non-purified fraction. The markers were expressed as fold-change to the non-purified fraction (NP) after normalization with hprt1 (n=3 mice per genotype). As shown in Fig 1B, the enrichment of mRNA for these markers was greater in ECs than in the non-purified fraction (NP). The immune system marker ptprc, and muscular cell marker alpha-SMA were higher enrichment in the NP fraction. Finally, the purity of ECs was further determined after several days in culture using the immunofluorescence of VE-cadherin that confirmed the homogeneity of the isolated cells (Fig 1C).

**Figure 1:**
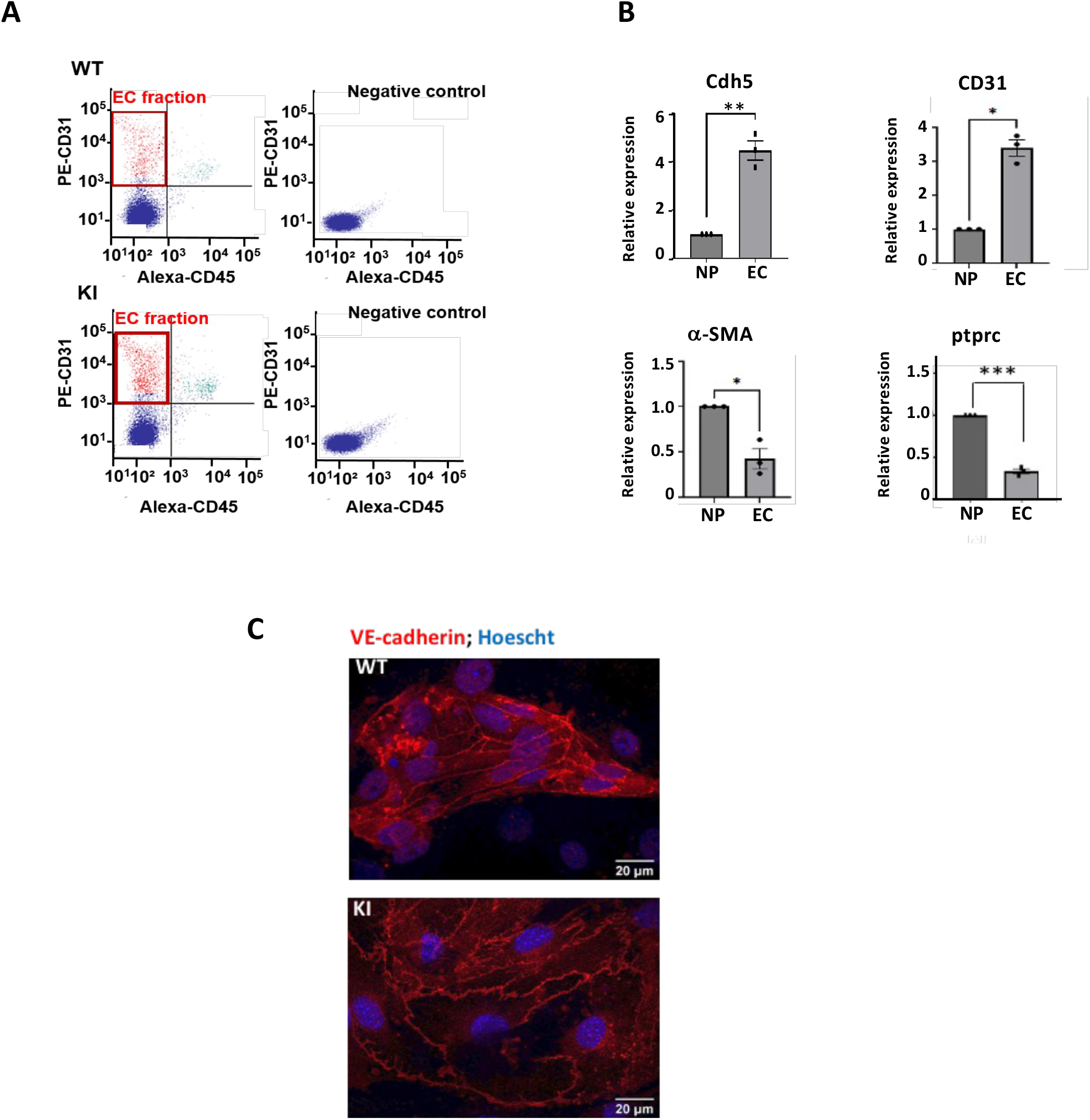
Characterization of Lung Endothelial Cells (ECs) from mutant (KI) and wild type (WT) mouse. **A** Representative sorting of EC using fluorescent activated cell sorter (FACS). Mouse cells were stained with PE-CD31 and Alexa488-CD45. The ECs population are represented in red (88.3%), the immune cells (0.02%) and the other cell types (11.40%) are in blue. **B**. Representative levels of several endothelial markers validate the purity of the isolated ECs. Total RNA was reverse-transcribed and amplified using exponential non saturating conditions with specific oligonucleotides for Cdh5, and CD31. PCR products were separated on ethidium bromide-stained 2.5% agarose gel. α-SMA, and ptprc were used for non-purified fraction. The markers were expressed according fold-change to the non-purified fraction (NP) after normalization with hprt1. (n=3 mice per genotype). Statistical significance was calculated by Student’s t-test (*p-value<0.05). **C**. Immunofluorescence of VE-cadherin in the isolated ECs. Nuclei were stained with Hoescht (blue). The slides were imaged with the Axio Imager M2 Zeiss microscope (magnification X63).

### Isolated ECs from mutant mouse have modified properties of migration, adhesion and proliferation

In most adult tissues, ECs are in quiescent state, which is a reversible state, and ECs can enter the cell cycle and expand to meet physiological conditions. In the lung, this is not the case as this tissue is not submitted to hormones or growth factors for its physiological functions. We were wondering whether the ECs from mutated mice would exhibit different properties involved in the angiogenic processes. Vascular ECs are critical to this process, since angiogenesis occurs via migration of ECs from pre-existing vessels, endothelial cell movement and proliferation (26). To test this hypothesis, a comparative study was carried out between the two genotypes for the properties of EC migration, adhesion and proliferation.

We first investigated the impact of mutated VE-cadherin on EC migration as it is one of EC functions which involves the dynamic and the interaction between ECs and VE-cadherin (27). The cell culture wound closure assay is one of the basic readouts for characterizing the migratory activity of cells (28). It is a measure of the lateral 2D migration of EC in cell culture. The isolated primary ECs from lung were submitted to wound closure assay using the WoundMaker to create 96 homogeneous scratches in each microplate well. The wound healing was visualized in real time using the IncuCyte ZOOM™ and imaged over 8 h time period. As a result, the analysis of the lateral scratch wound showed a significant difference in the rate of migratory response between ECs from KI (22.45% ± 5.207) compared to WT (13.24% ± 5.17) (p-value=0.034) after 20 hours suggesting that the mutant ECs migrate slower than the WT ones (Fig 2A,B).

**Figure 2:**
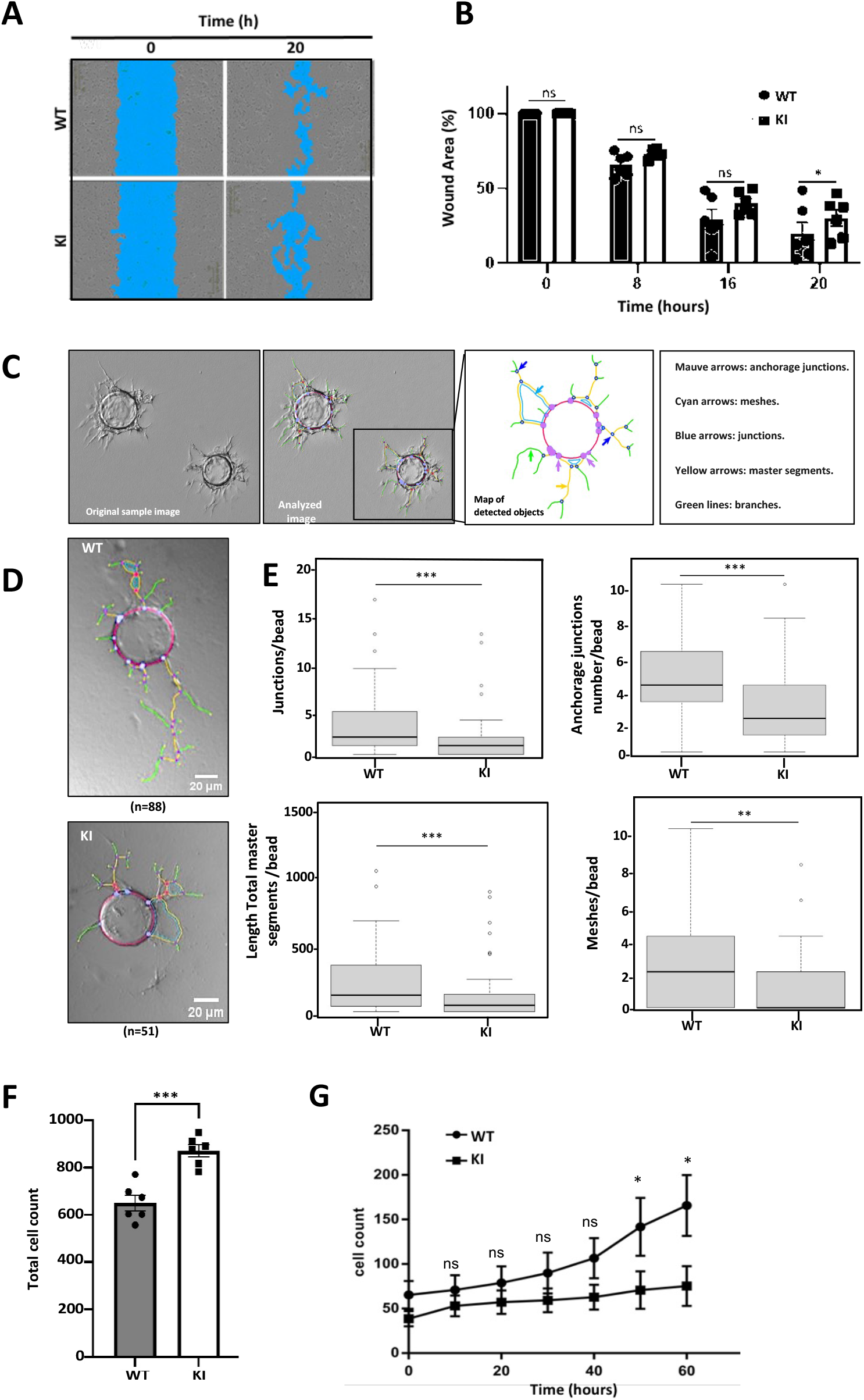
Lung ECs from mutated VE-cadherin mice have modified migration, adhesion and proliferation properties. **A**. Cell culture wound closure assay of ECs from WT and KI. ECs were seeded onto 96-well microplates to reach confluency within 48h. The scratch was made with the Incucyte® 96-Well Woundmaker Tool. The blue region denotes the scratch wound mask over time (t=0, t=20 hours). **B.** The scratch closure was monitored and imaged in 8 h intervals. The percentage of gap area was followed by repeat scanning in the live-cell analysis software until 20h. Magnification (X10) (n=3 mice per genotype). **C**. 3D Fibrin gel beads in vitro assay. Representative images of EC coating beads sprouting after 4 days. Quantification of morphometrical parameters of the capillary network was performed by a computerized method (see on the bottom panel) on images taken on day 4. **E.** Representative parameters measured were number of anchorage per junctions, number of junctions, total master segments and number of meshes per beads. 88 beads for WT and 51 for KI were quantified**. F** Cell adhesion assay. Cells were seeded at 1500 cells/96 wells onto fibronectin coated plate and incubated for 30 min and stained with Hoescht. The nuclei were counted for each well (n=6). **G**. Proliferation assay for WT and KI cells. ECs were seeded at a density of 1000 cells/mL onto a 96 well plate (n=3 mice per genotype). Results are expressed as the mean of triplicate determinations and the experiment was repeated three times with similar results. All bar graphs showed mean ± SEM. Statistical significance was calculated by Student’s t test two-tailed unpaired t-test, *p-value<0.05, ** p-value<0.01, **** p-value<0.0001 (*<0.05).

We next studied *in vitro* assays for sprouting angiogenesis properties in which isolated ECs WT and KI coated on beads were allowed to sprout in fibrin gel. After 4 days, KI ECs and WT ECs elicited different angiogenic responses (Fig 2C). Thanks to a powerful automatic quantification method, we were able to determine the changes in the morphometrical parameters of the neovessels formed in the fibrin matrix (22). Parameters chosen to our study were: number of junctions per bead which is a sign of pseudo-vessels complexity, number of anchorage junctions per bead, assimilated to the sprouting capacity, total master segment length per bead, associated with the elongation, the development of the pseudo-capillaries capacity, and number of meshes per bead for the capacity to establish mesh network. KI ECs exhibited less angiogenic process than WT ECs, as the number of junctions per bead, the number of anchorage per junctions per bead, the total master segment length per bead and the number of meshes per bead were significantly decreased compared to the WT ECs (Table 2).

**Tab 1.**
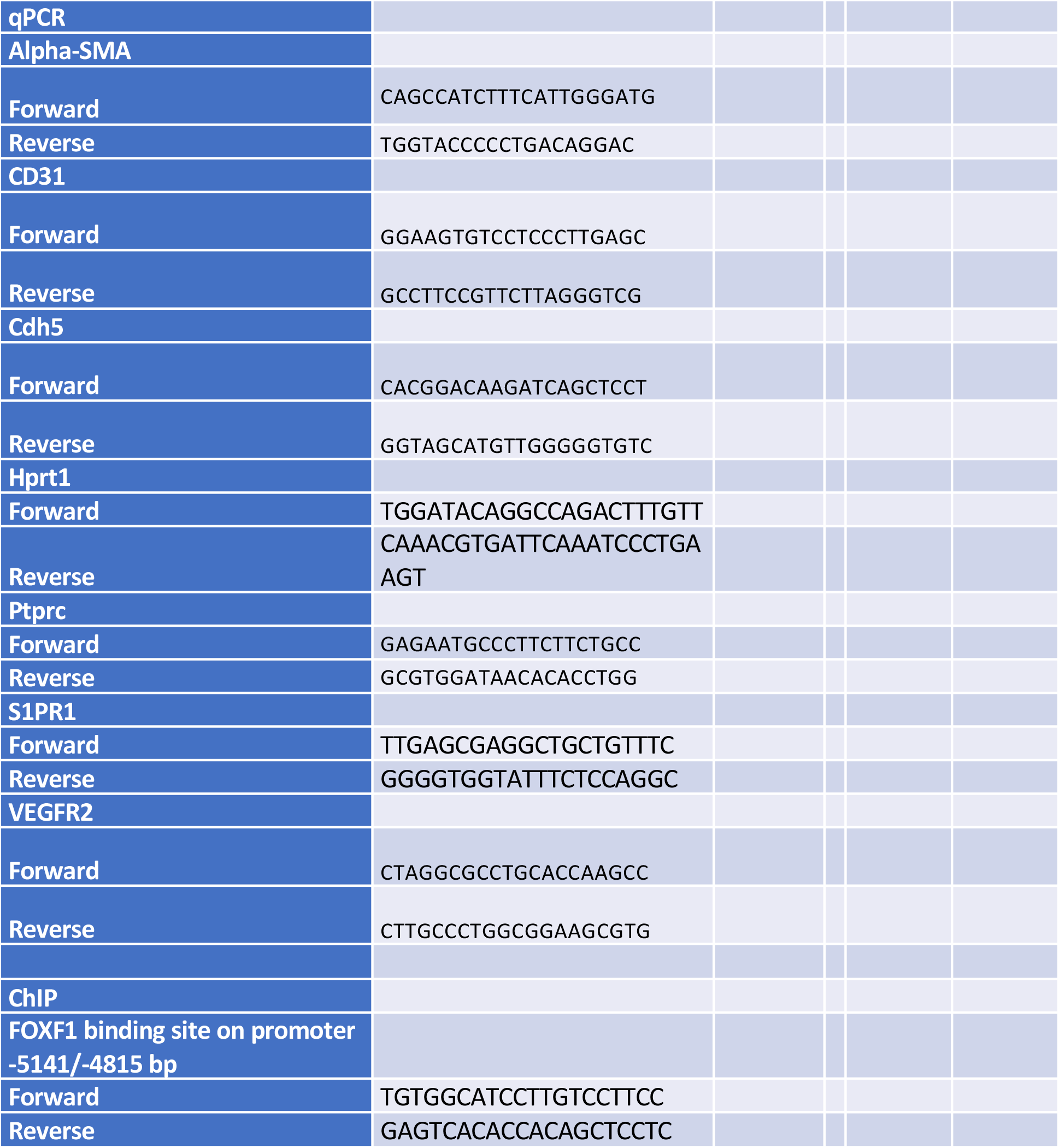
List of primers sequence in mouse.

**Table 2:**
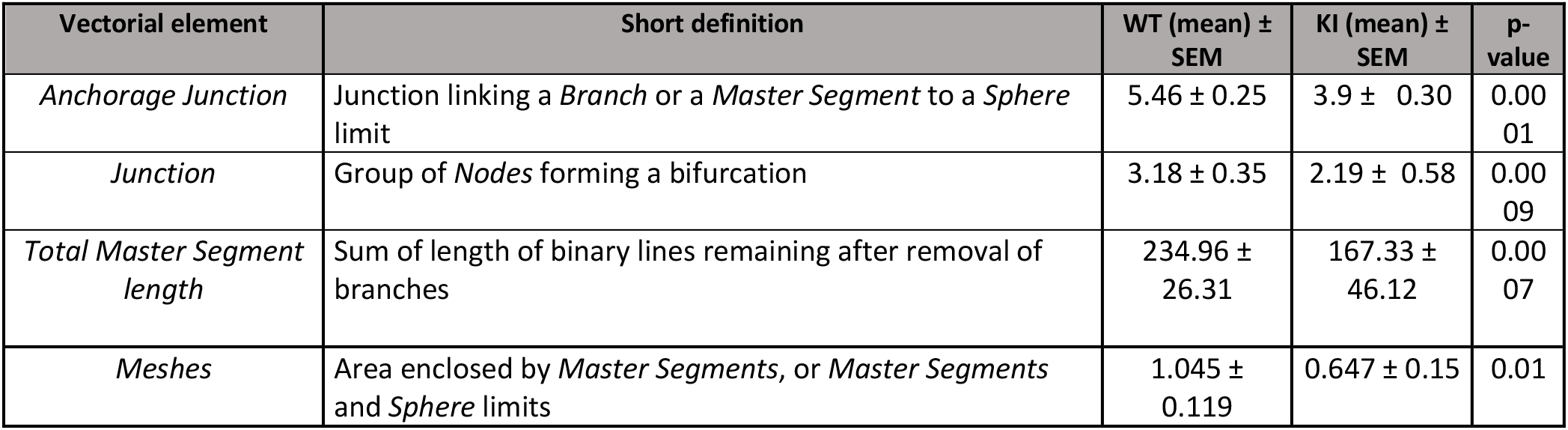
Vectorial objects characterized and quantified by the software analysis for the WT and KI cellular trees.

Altogether, these results found with the wound assay and the fibrin beads assay strongly suggest that the ECs from lungs of the KI mice present slightly difference in their migratory properties.

Next, because the rate limiting step for ECs to start to form a monolayer from blood vessel is the initial adhesion to the extracellular matrix, we wanted to determine if any difference was found between the two genotypes (29). Thus, we used fibronectin which is the most classical exogenous substrate for ECs as it represents the classical adhesion-promoting protein. Briefly, the ECs were seeded (1500 cells/well) onto fibronectin-coated plates and incubated for 30 min at 37°C. At the end of the incubation time, the nuclei were staining with Hoescht and counted (30). As a result, the interaction with fibronectin was significantly higher (37.625 ± 9.23) in KI than in WT (26.8 ± 3.258, p-value=0.012) (Fig 2D). This data suggest that the ECs from KI mice formed the first few bonds during initial cell-substrate contact much faster than the ECs from WT mice.

We next wondered whether VE-cadherin mutant could affect EC proliferation as VE-cadherin was described to act as a negative regulator of EC proliferation (31,32). To test this hypothesis, the ECs from both genotypes were seeded in 96 well-plates and the proliferation was followed for up to 60h. As a result, the ECs from WT mice started to proliferate between 40 to 60H, while the ECs from the mutant mice remained at a constant rate over the time period tested (WT 141.85 ± 32.52 vs KI 70.66 ± 14.86, p=0.040 for 50h; WT 165.71 ± 34.14 vs KI 75.22 ± 24.25, p=0.037 for 60h) (Fig 2F). This data is consistent with our previous results showing that ECs from mutant mice are more adherent to the external substrate. Altogether, these data suggest that the population of ECs isolated from mutated mice showed an increased adhesion capacity and impaired migration and proliferation properties.

### Transcriptome of ECs from lungs of WT and KI mice

Lung-specific EC harvests yielded high quality RNA for gene expression analysis (data not shown). The RNA extracted from the cells was subjected to Next Generation Sequencing analysis (RNA-seq), and the resulting reads were aligned along 94,818 transcripts of the mouse genome (Fig 3A). The global transcriptional changes across the two groups was visualized by a Volcano plot where the log fold change of each gene is represented (x-axis) in function of (−log10) of its adjusted p-value (y-axis). Genes with a corrected p-value less than 0.05 and a absolute log2(Fold change) > 1 are indicated by red dots. 884 genes were differentially expressed between ECs from KI mice and WT with 766 genes downregulated and 118 genes upregulated in ECs from KI mice. The top 30 differentially expressed genes based on the corrected p-values sorted by the averaged expression value across samples are illustrated in Fig 3B. The reproducibility among biological triplicates demonstrate the fidelity of the identification, isolation, and profiling of lung-specific ECs. One of the gene that is the most differentially expressed in the ECs from KI mice is the Sphingosine-1-phosphate-receptor 1 (S1PR1). Differentially expressed genes were subjected to the GO enrichment analyses to extract biological knowledge (Fig 3C). Over the twenty ontologies analyzed, the most significant were related to cell adhesion and angiogenesis included the vascular development and blood vessel morphogenesis which are the most deregulated functions linked to the endothelial characteristics. Several examples of genes were found to be downregulated in the KI such as*, Apln* (encodes for apelin), *Slit2* (encodes for Slit homolog 2 protein)*, Robo4* (for Roundabout homolog 4), *Egfl7* (encodes Epidermal growth factor-like protein 7), *Mmrn2* (encodes for Multimerin-2)*, Ccbe1* (for Collagen and calcium-binding EGF domain-containing protein 1), *Kdr* (encodes for VEGFR2), *Ecscr* (for Endothelial cell-specific chemotaxis regulator), *Mmp2* (for typeIV collagenase). These genes were involved in the positive balance of angiogenesis and were down regulated in the KI mice (33–41). Among the other family of genes involved in the cell adhesion, several were known to influence EC-matrix interaction, including *Igsf5* (for Immunoglobulin superfamily member 5) (42,43), *Vcam1* (for Vascular cell adhesion protein 1)(44), *Jam2* (for Junctional adhesion molecule B) (45), *Lamb1* (laminin β1)(46). These genes were downregulated in the KI mice and the inhibition of their expression resulted in a greater cell attachment to ECM. In contrast, the genes *Jag1* (Jagged 1) (47), *Spock2* (sparc/osteonectin, cwcv and kazal-like domains proteoglycan 2) (48) and *Itga5* (Integrin Subunit Alpha)(49) were upregulated in the KI. Another interesting function affected by the mutation of VE-cadherin is the vascular permeability since the site Y685 plays a key role in this process. The most important genes that were up-regulated in KI mice and related to permeability were *Slit2*, *Bmp6* (bone morphogenetic protein 6) (50, 51), *Ramp2* (receptor activity modifying protein 2) (52) while *Plec* (plectin) (53) was a gene known to be involved in the integrity and found to be upregulated. In the KI mice.

**Figure 3:**
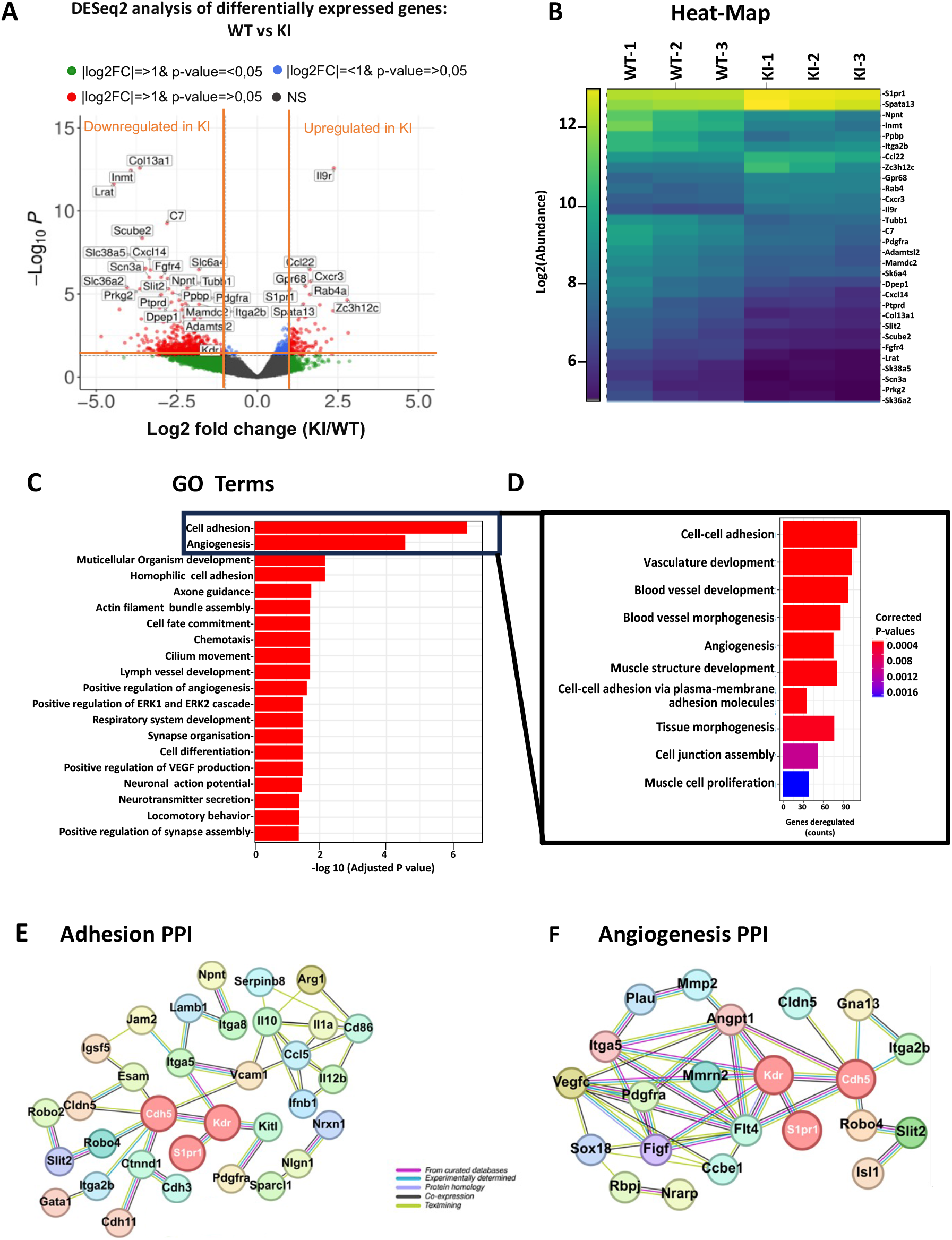
Gene expression profiling of FAC-sorted lung ECs (mutated versus WT) showed genes involved in the cell adhesion and angiogenesis. **A.** The global transcriptional change across the groups compared was visualized by a Volcano plot. Each data point in the scatter plot represents a gene. The log fold change of each gene is represented on the x-axis and the log1 of its adjusted p-value on the y-axis. Genes with an adjusted p-value less than 0.05 and a log2 fold change greater than 1 are indicated by red dots. These are up-regulated genes. Genes with an ajusted p-value less than 0.05 and a log2 fold change less than −1 are indicated by green dots. These represent down-regulated genes. **B.** A bi-clustering **h**eatmap was used to visualize the expression profile of the top 30 differentially expressed genes sorted by their adjusted p-value by plotting their log2 transformed expression values in samples. **C.** Gene Ontology terms of all genes. **D.** Gene ontology terms classified according to their function.. **E**. Deregulated gene-gene interaction network analysis involved in cell–cell adhesion. **F.** Deregulated protein-protein interaction network of DEGs involved in angiogenesis. Analysis was performed using String software. Cdh5 is encircled. The thickness of the line between two genes is correlated with the strengh of data support.

The gene ontology database revealed the top ten biological processes (BPs) from the enrichement of specific significant genes (corrected p-value < 0.05 and absolute log2(Fold change) > 1) (Fig 3D). Using the STRING database to retrieve interactions between DEGs identified from RNAseq, a network of interacting proteins/genes was established to examine cluster coalescence (Fig 3E,F). Cluster analysis revealed that the largest protein-protein interaction (PPI) was in the process of adhesion and angiogenesis. S1PR1 and KDR were found to interact with Cdh5 in cell-cell adhesion and angiogenesis protein-protein networks (Fig 2D, 2E). These data suggested that the gene expressions encoded by VE-cadherin mutation might be involved in the main endothelial functions such as cell-adhesion and angiogenesis.

### S1pr1 is a direct transcriptional target of the FOXF1 in lung ECs from KI mice compared to WT and maintains VE-cadherin at the plasma membrane

Our transcriptomic data pointed to S1PR1 as the gene specifically upregulated in the mutant mice when compared to the WT. Therefore, we wondered whether the transcription factor forkhead box F1 (FOXF1) which was very abundant in quiescent adult lungs was involved in this process (54). In addition, FOXF1 deficiency was associated with a decreased S1PR1 expression in vivo and in vitro (55,56). Therefore, we analyzed whether FOXF1 directly regulate *S1pr1 promoter* in both mice genotypes as the transcriptomic analysis showed that S1PR1 was one of the gene upregulated in the KI ECs with the higher adjusted p-value (p>10^-12^). There are four potential FOXF1 binding sites that were identified in the −6.0-kb promoter region of the mouse S1PR1 gene. It was shown in the literature that FOXF1 specifically bind to the −3880/−3876–base pair (bp) and −5141/−4815-bp regions of the *S1pr1* promoter (54) (Fig 4A). We thus performed a chromatin immunoprecipitation assay (CHIP) using mouse lung nuclei extracts from KI and WT to determine whether FOXF1 protein could physically bind to S1pr1 promoter. We chosen the −5141/−4815-bp DNA fragment to perform RT-qPCR with primers (see table S1) designed to capture the FOXF1-binding site near the transcription start site of the *S1pr1* promoter. EconoTaq Plus Green according the following cycles: 3 minutes at 95°C, 34 cycles: 95°C 15 seconds; 62°C 15 seconds; 72°C 20 seconds, 72°C 10 minutes. The DNA concentration was measured and the same amount of DNA (400ng) were loaded on the agarose gel for analysis. FOXF1 cross-linked protein/DNA complexes were immunoprecipitated with rabbit anti-FOXF1 (Fig 4B). The relative fold change of FOXF1 binding to S1PR1 promoter between WT and KI lungs were analyzed in 3 mice per genotype. We have clearly identified that the FOXF1 protein directly bound to promoter DNA of *S1pr1* in the both genotypes with a much higher quantity in KI than in WT (fold-change of 7 ± 1.76, p-value=0.023) (Fig 4C). The specificity of the binding of FOXF1 was further confirmed by Western blot where FOXF1 was only present in the CHIP samples and absent in the supernatant (data not shown). These finding suggest a direct transcriptional regulation of S1pr1 by FOXF1 in EC from mutant mice. Analysis of the S1PR1 RNA and protein were further performed. Subsequent real-time PCR confirmed the above findings and showed a significant induction of S1PR1 mRNA expression, with an average fold change of 3.26 ± 0.310 (p-value=0.011) (Fig 4A). When normalized to the level of β-actin, immunoblotting analysis of lung protein extracts with the antibody against S1PR1 showed a four-fold increased (p-value=0.034) of the immunoreactive protein in the KI mice compared to WT.

**Figure 4:**
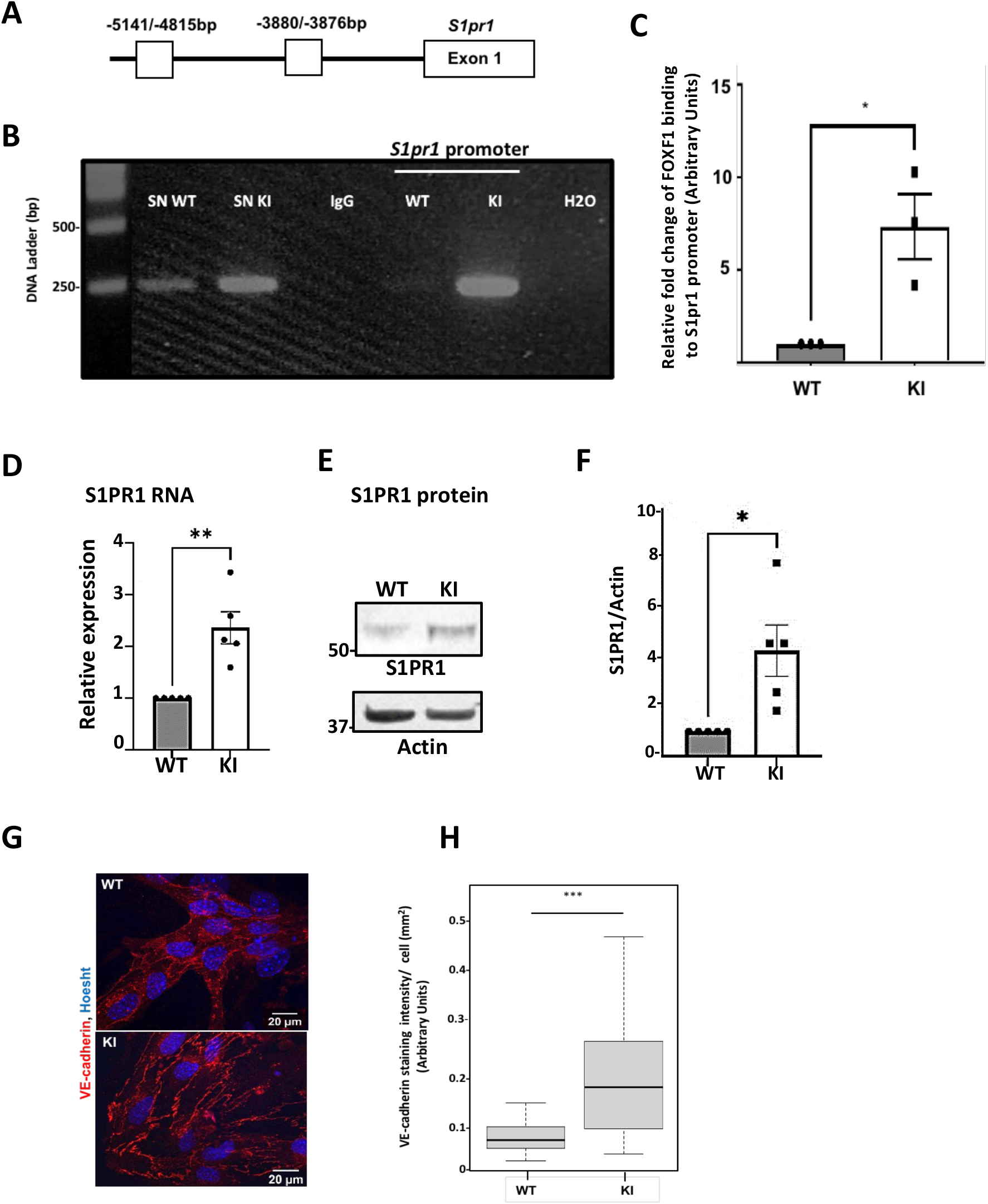
FOXF1 directly binds to the *S1PR1* promoter in KI mice vs WT and increase the production of S1PR1 at the RNA level and the protein level. **A.** Schematic drawing of the mouse S1pr1 promoter region with potential FOXF1 DNA binding sites (white boxes). FOXF1 protein binds to the −5141/−4815-bp and −3880/−3876-bp of S1pr1 promoter regions**. B.** FOXF1 cross-linked protein/DNA complexes were immunoprecipitated with rabbit anti-FOXF1. The same quantity of DNA was loaded onto the agarose gel for each fraction (Supernatant (SN) and S1PR1 promoter regions bound to FOXF1). RT-qPCR was performed with primers designed to capture the FOXF1-binding site near the transcription start site of the *S1PR1* promoter, negatively and positively controlled by IgG and total input without immunoprecipitation. **C**. The relative fold change of the binding of FOXF1 on S1PR1 promoter regions between WT and KI (n=3 mice per genotype) was quantified using Image-J software (NIH, Bethesda, MD). Molecular mass standards (in kDa) are shown at left. **D,E E.** Representative immunofluorescence of the VE-cadherin on isolated EC. Isolated ECs from lungs were cultured for 4 days and then fixed with PFA and stained with the VE-cadherin antibody. Hoescht was used to stain cell nuclei. Magnification (X63). **F**. The intensity of VE-cadherin staining at membrane was quantified. Images are representative of 3 mice per genotype. (n=29 for the WT and n=17 for the KI). Statistical analysis was performed using Mann-Whitney test (***p-value<0.001) as described in Methods. All analysis of the relative expression of the proteins was measured by densitometry of autoradiographs using Image-J software (NIH, Bethesda, MD). All bar graphs were mean ± SEM. Statistical analysis were performed using Student t-test (*p-value<0.05).

As S1PR1 was shown to maintain VE-cadherin at the membrane of ECs (54), VE-cadherin assembly into cell-cell junctions was then investigated in lung ECs from WT and KI mice in culture. The isolated ECs were stained with the VE-cadherin antibody and the immunofluorescence found at the plasma level was analyzed. The VE-cadherin area at the membrane and the intensity of the immunostaining per cell were reported for the two conditions. The VE-cadherin fluorescence intensity observed in the KI ECs was higher than that of WT ECs (100 ± 6.52 for WT vs 189.7 ± 21.06 for KI (p-value 0.0001) (Fig 4D,E). Thus FOXF1 induces endothelial barrier function through transcriptional induction of the S1pr1 promoter, which, in turn, cause activation of S1PR1 signaling and stabilization of VE-cadherin at the plasma membrane in the KI model.

### Mutated VE-cadherin downregulates VEGFR2

Previous studies showed that the loss of *S1PR1* in ECs induced vessel hyper-sprouting in the retina due to increased VEGFR2 activity (57). Therefore, we wondered whether the expression of VEGFR2 both at transcriptional and at the protein level was modified. As a result, the relative expression of VEGFR2 mRNA was found to be decrease by 50% in the KI mouse vs the WT (p-value= 0.012). When normalized to actin, the protein expression of VEGFR2 was also decreased (p-value=0.02) (Fig 5A) as well as the phosphorylation of VEGFR2 at site Y^1173^ (KI: 0.53 ± 0.23 vs WT 1.71± 0.10; p–value= 0.009). In agreement with RT-PCR experiments and protein analysis, this data confirmed that VEGFR2 was less active in the KI mice than in WT (Fig 5B). In regards of these results, we wanted to verify whether this VEGFR2 modification was related to the VEGFA level. Protein analysis was performed by Western blotting by use of an antibody that recognized the isoform 165 as a 23-kDa protein. No commercial antibody was available to detect specifically the 188 isoform. Thus, in our hands, we found a 23-kDa protein as well as the 46 kDa present in the same amount in the lung extracts of the two genotypes (Fig 5C). Altogether, these results suggested that VEGFR2 was inhibited both at a transcriptional level and a protein level without any change in VEGF expression.

**Figure 5:**
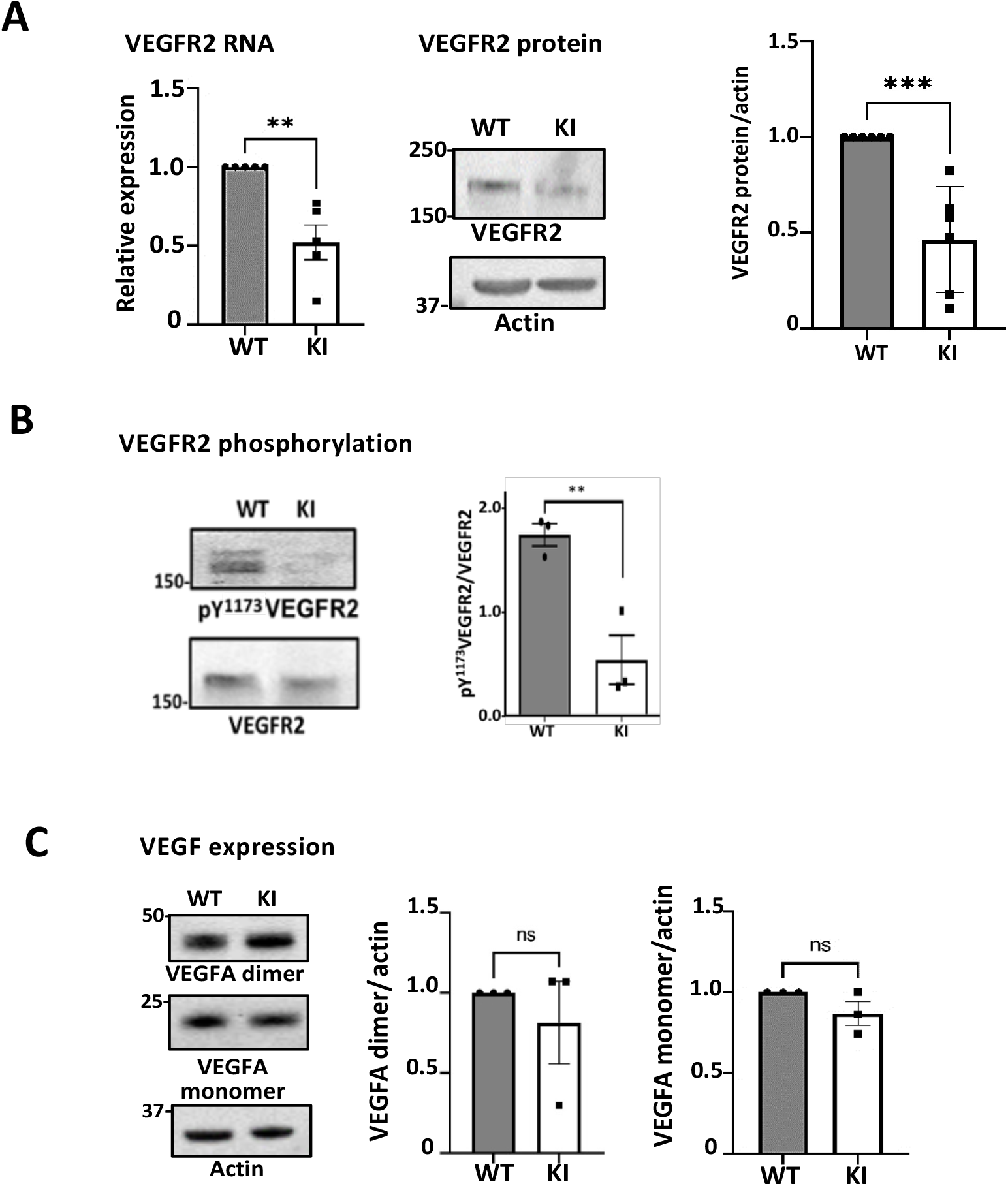
Effect of VE-cadherin mutation on, VEGFR2 and VEGF expression. **A**. Expression of S1PR1 increased in the KI mice. S1PR1 RNA expression level from isolated endothelial cells was quantified as the fold-change of WT after normalisation with hprt1. Immunoblotting analysis of lung protein extracts (30 μg) was performed using the antibody against S1PR1 (n=5 mice per genotype) and normalized with actin. **B**. Expression of VEGFR2. RT-PCR analysis was performed from isolated EC RNA extract. VEGFR2 RNA expression level was expressed as the fold-change of WT after normalisation with hprt1. Immunoblots was performed from lung protein extract (30µg) and protein level was normalized with actin (n=5 mice per genotype). **C**. VEGFR2 phosphorylation was analyzed using the anti pY^1175^ antibody (30μg). **D**. Analysis of VEGFA dimer and monomer by Western blot (n=3 mice per genotype). All inmmunoblots were carried with 30 μg of total proteins and protein level was normalized with actin. Molecular mass standards (in kDa) are shown at left. All analysis of the relative expression of the protein were measured by densitometry of autoradiographs using Image-J software (NIH, Bethesda, MD). Statistical analysis performed by Student’s t-test, (***, p-value<0.001. (*p-value<0.05). Non significant ns)

### VE-cadherin mutant is phosphorylated at site Y^731^ and is associated with active Src but to a lesser extent with β-catenin

To get further insights into the mechanisms of action of the mutated form of VE-cadherin on the possible signaling pathways, we firstly determined the level As it was reported in the literature that the other cytoplasmic VE-cadherin sites (Y^658^ and Y^731^) were also substrate for Src kinase activity, we wondered whether the VE-cadherin mutant was phosphorylated on these sites (60). Lung proteins extracts were tested with the antipY^658^ VE-cadherin antibody and were normalized to the level of β-actin expression. While only a small difference was detectable for the site pY^658^VE-cadherin between KI and WT (KI 0.38 ± 0.059 vs WT 0.553 ± 0.2; p-value= 0.45) (Fig 6A), the phosphorylation of the site Y^731^ was more prominent in the KI than in the WT (KI 1.48 ± 0.14 vs WT 0.82± 0.13, p-value=0.041) (Fig 6B). This results demonstrate that the mutated VE-cadherin is naturally more phosphorylated in lungs *in vivo* as compared to WT.

**Figure 6:**
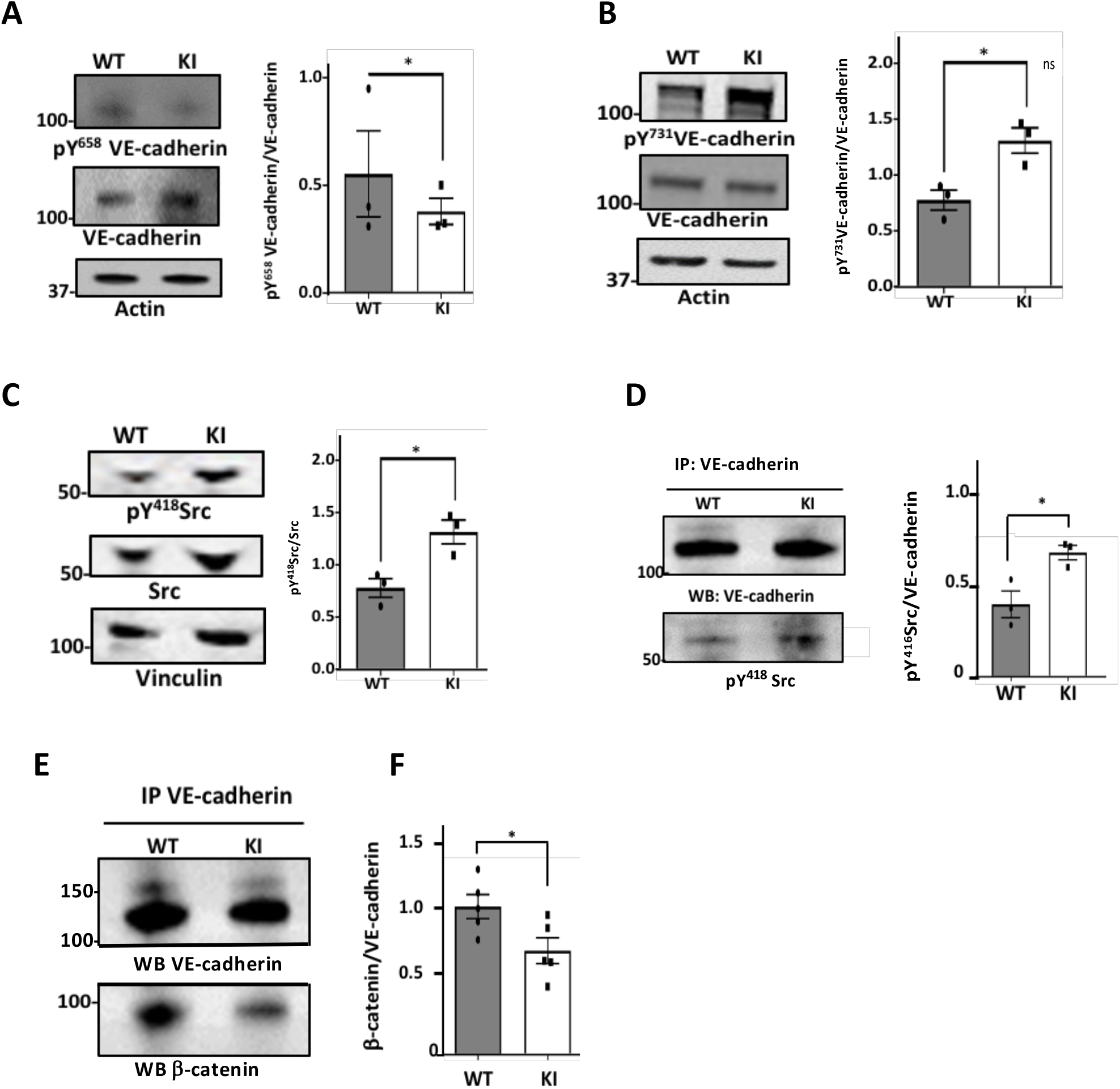
VE-cadherin mutant is phosphorylated at site Y^731^ and is associated with active Src but to a lesser extent with β-catenin. **A.** Western Blot analysis of pY^658^ VE-cadherin and VE-cadherin in WT and KI using 30 μg of protein lung extracts. Actin was used as loading control (n=3 independent experiments). **B.** Western Blot analysis of pY^731^VE-cadherin in WT and KI using 30 μg of lung extracts. Actin was used as loading control (n=3 mice per genotype). **C.** Western blot analysis of pY^418^ Src and Src were performed using 5 μg of protein extracts from the isolated ECs from WT and KI mice. Vinculin was used as loading control (n=3 mice per genotype). **D**. Representative western blot of VE-cadherin immunoprecipitate probed with pY^418^ Src antiboby (n=3 mice per genotype). **E**. Representative western blot of VE-cadherin immunoprecipitate from lung mice extract (100 μg) probed with β-catenin antibody (n=5 mice per genotype). Molecular mass standards (in kDa) are shown at left. Statistical analysis was performed using Mann-Whitney test (***p-value<0.001) as described in Methods. All analysis of the relative expression of the proteins was measured by densitometry of autoradiographs using Image-J software (NIH, Bethesda, MD). All bar graphs were mean ± SEM. Statistical analysis were performed using Student t-test (*p-value<0.05).

We next determined whether activated Src was found in ECs from WT and KI mice. To investigate this possibility, we performed immunoblot with the antipY^418^-Src antibody from ECs isolated from WT and KI mice. When normalized to vinculin, much to our surprise, activated c-Src (pY^418^) was highly increased in the ECs from KI versus WT (KI 1.29 ± 0.09 vs WT 0.77 ± 0.07, p-value= 0.02)(Fig 6C). These findings reveal that active Src can lead to the phosphorylation of the site Y^731^ of VE-cadherin which is consistent with the increased Src activity and Src-associated tyrosine phosphorylation events. Because the phosphotyrosine-containing proteins are binding sites for c-Src-homology-2 (SH2) domain, we wondered whether an association of VE-cadherin/Src could be observed *in vivo*. Lysates from lungs were subjected to immunoprecipitation with anti-VE-cadherin antibody and immunoblotted with anti-pY^418^Src antibody. It was found that Y^418^-Src signal was significantly prominent in VE-cadherin immunoprecipitates from the KI mice (0.60 ± 0.035; p-value=0.0269) and to a lesser extent in lung from WT (0.35 ± 0.0643) (Fig 6D). These findings confirmed that when VE-cadherin is phosphorylated, it is an anchor to maintain Src at the EC junctions, where it could exert its activity on junctional components. Previous published work have shown that a tyrosine-to-glutamic acid mutant Y^731^E-VE-cadherin impaired the binding of β-catenin without displaying barrier function (61). Because the phosphorylation at site Y^731^ was higher in VE-cadherin mutant than in WT, therefore, we wondered whether an association β-catenin/VE-cadherin mutant was influenced by the status of phosphorylation of VE-cadherin *in vivo*. Lysates from WT and KI mice were thus subjected to immunoprecipitation with VE-cadherin antibody and immunoblotted with the antibody to β-catenin. While the presence of β-catenin was detected in each immunoprecipitate, a significant decrease in the interaction between VE-cadherin and β-catenin was found in KI mice (0.798 ± 0.098) vs WT (1.136 ± 0.092; p-value=0.0369) (Fig 5E). In agreement with Potter et al (61), these data confirmed that some β-catenin molecules were removed from the pY^731^VE-cadherin complex of adherens junctions.

### EC mutation induced vascular alterations of lungs

Based on our transcriptomic data from ECs derived from lung mice, we wondered whether the mutated VE-cadherin could affect the morphology of lung vessels. Pulmonary vasculature was analyzed from lung sections (Figure 7A). The total vessel number was expressed as number of blood vessels per lung section for each group. The results demonstrated a strong decrease, in KI mice n=287 as compared to WT (n=411) (p-value=0.036) (Fig 7B). Assessment of pulmonary vascular thickness was made by comparing pulmonary arteries of similar diameter (75-150µm) on each lung section. The medial wall was attested by the measurement of the standard medial thickness (WT 26.87 ± 1.08; n=55 versus KI 41.39± 6.48; n=81 respectively, p-value=0.049) (Figure 7C) which was the ratio of medial area to total area (WT 0.58 ± 0.01 versus KL 0.66 ± 0.01 respectively, p-value=0.0000044). We assessed also the lumen area which was significantly narrower in the KI mice (WT 5550.23 μm2 ± 203 versus KI 4864.7 μm2 ± 280; p-value= 0.0098). An increase of 2.06 fold in the ratio of medial area to luminal area was also noticed to be significant in the KI mice (WT 1.49 ± 0.065 versus KI 2.99 ± 0.506, p-value=0.000013).Therefore, the morphometric analysis of the lung suggested a vascular remodeling in the KI.

**Figure 7.**
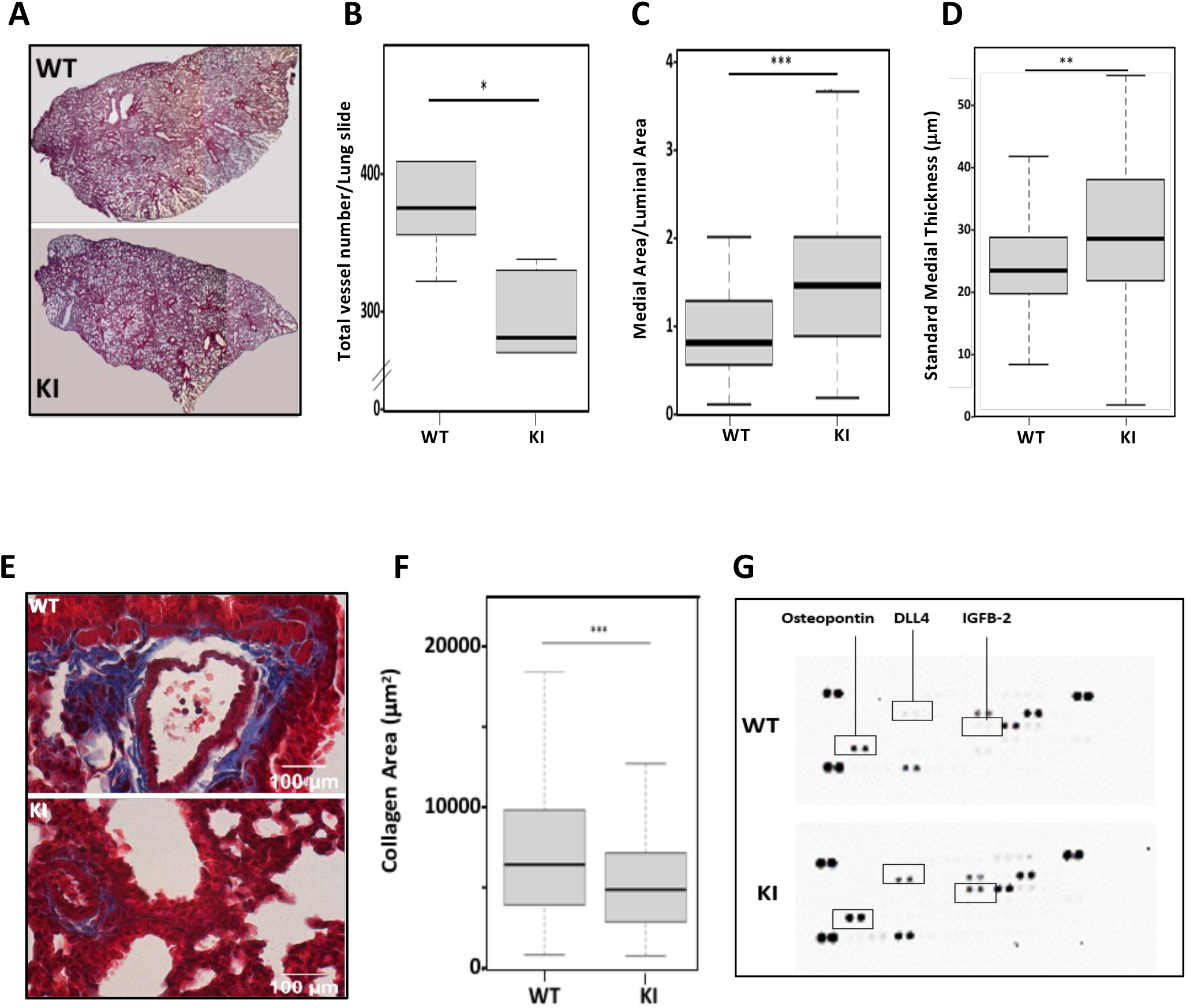
*In vivo* analysis of histological and morphometric evidence of thickened pulmonary arteries in the mutant and WT mouse: **A.** H&E from lung paraffin sections were used to count small pulmonary arteries of comparable diameter (75-150 μm). Images are representative of 4 mice per genotype**. B.** Medial area over luminal area ratio shows an increase in KI mice. **C.** Morphometric evidence of thickened, remodeled small pulmonary arteries. Standard medial thickness was calculated as the ratio of medial area to inner circumference. **D**. Representative histological profiles of paraffin-embedded lung sections stained haematoxylin-eosin and Masson’s trichrome in WT and KI showing marked medial thickening of smaller pulmonary arteries from KI mouse vs WT mouse. Magnification (x40), scale bar 100µm. Images are representative of 4 mice per genotype. **E,F.** Collagen fibrils stained with Masson trichrome’s were quantified using Image-J software (NIH, Bethesda, MD). Statistical analysis was performed using Mann-Whitney test (***p-value<0.001). **G**. « Mouse Angionesis Array » in the BAL fluid was performed according to the manufacturer’s instructions. The same amount of proteins was applied onto the membranes. The relative intensity of the array spots between the two genotypes were quantified using Image-J software (NIH, Bethesda, MD). (n=4 mice per genotype). All bar graphs showed mean ± SEM.

As collagen deposit was an indicator of fibrosis, we assessed whether any deposition of collagen occurred. Immunohistochemistry was performed with Harris’s hematoxylin-eosin and trichrome Masson staining on lung slides. Collagen fibrils from Masson trichrome’s staining were quantified using Image J software. The morphometric analysis demonstrated a significant reduction in collagen fibers around the walls of small pulmonary arteries from KI lung (mean 70039.02 ± 3142) as compared to WT (mean 53206 ± 3383.2852) (p-value 0.0008) (Fig 7D,E).

We wanted then to determine the functional properties of the pulmonary endothelium which forms a tight barrier and actively regulates paracellular extravasation of proteins, solutes and fluids to control interstitial fluid homeostasis. A bronchoalveolar lavage (BAL) is an important and most commonly used technique to study inflammatory cell infiltration, biochemical, and molecular changes. BAL analysis was performed in WT and KI mice to determine the subtle protein expression using a “Mouse Angiogenesis antibody Array” which is sensitive technique. Among 53 immunoreactive proteins that could be detected with this array, only five proteins were found in BAL from WT while ten proteins were in KI mice. Among them, three cytokines exibited a significant increase in the KI such as DLL4 (fold increase 2.38), insulin-like growth factors 2 (IGFB2) (fold increase 1.31) and osteopontin (fold increase 1.58) (Fig 7F). Altogether, these data suggested that the mutated VE-cadherin presented vascular remodeling and no fibrosis induction, and an increase of proteins known to have negative effect on angiogenesis.

## DISCUSSION

Our data demonstrate for the first time that a single point mutation in VE-cadherin cytoplasmic domain dysregulated endothelial gene expressions in lung ECs. Of importance, this mutation was obtained because it was Src kinase site for which the sequence LY^685^AQV fits the YxxV/I/L motif, which was found to be a consensus sequence for Src family kinase (12,13). Here we addressed the issue of the role of this mutation in EC gene expressions because there are several VE-cadherin partners including β-catenin whose interaction could be dependent on the phosphorylation status of VE-cadherin. As such, β-catenin may translocate to nucleus and bind to regulators of gene transcriptions (63).

## PROPERTIES OF ECs

First of all, we attempted to isolate ECs from lung as it was demonstrated that VE-cadherin expression in the lung was greater than in other tissues (11) and of importance because it was shown an endothelial heterogeneity along the vascular tree of different organs (1, 64). The vascularization of the lungs is mostly composed of capillary ECs which is a significant component of the alveolar-capillary membrane, a thin tissue barrier responsible for gas exchange between alveolar air and the pulmonary circulation (65). ECs from both mice exhibited a high level of VE-cadherin as determined by RT-PCR and immunofluorescence which is in agreement with our previously published data indicating that in KI animals VE-cadherin was expressed to the same extent than in WT animals (15). To further characterize the ECs from lung, we determined their angiogenic functions. We demonstrated that the EC from KI had a slower rate of migration, both in the wound and in the fibrin beads assay, but they exhibited increased adhesion properties to extracellular matrix likely through integrin receptors (29). Cadherin-mediated adhesion also directly initiates both mechanical and biochemical signals known to affect focal adhesion formation. The focal adhesion kinase (FAK) is a tyrosine kinase that is involved in the assembly of focal adhesions and thereby promotes cell spreading on the extracellular matrix and its autophosphorylation site reflects its activity (66). Whether or not FAK is phosphorylated at site Y^397^ is still under investigation. Altogether, our data demonstrate several differences in angiogenic properties of the ECs of the two genotypes.

## TRANSCRIPTOMIC ANALYSIS

To obtain a deeper understanding of the mutation of VE-cadherin on the endothelium, we took the opportunity of this mouse model to perform for the first time a transcriptomic study of lung endothelial cells from 4 weeks-old male mice which can be consider as adult mice. Diversity in ECs function has been recognized for decades. However, we could analyse the vascular endothelial phenotypes by comparison of the two genotypes. The global transcriptional change across the groups compared was visualized by a Volcano plot. It is interesting to note that more than 800 genes were modified in this mouse model with 766 genes downregulated and 118 genes upregulated in ECs from KI mice. It shows the importance of VE-cadherin tyrosine phosphorylation on the site 685 in normal physiological conditions. Further studies will be needed to understand how these gene expressions can be modified “only” by the status of phosphorylation of one tyrosine.

The most specific endothelial transcription program that we found was related to cell adhesion and angiogenesis. Our data were coherent with gene expression profiles which revealed a panel of differentially expressed cell adhesion molecules that contribute to enhanced cell–cell adhesion in microvascular cells (71). Several evidences have established that the pulmonary microvascular endothelium is much less permeable than that of capillaries in other organs (72,73). Cultured rat pulmonary microvascular ECs demonstrate reduced basal and inflammatory-mediated permeability to solutes. The retention of site-specific differences in permeability in multiply passaged ECs might explain these differences. Pulmonary micro- and macrovascular ECs demonstrate significant cell type–specific differences in DNA methylation pattern. In contrast, ECs in the bronchial vasculature are more leaky, are more responsive to inflammation, and have a far greater capacity for angiogenesis compared with ECs from the pulmonary vasculature (72).

## FOXF1-S1PR1-VE CAD MEMB

Within the sets of genes upregulated by the mutated VE-cadherin, we selected S1PR1 because it has been well described in playing an important role in the barrier integrity (74). Moreover, S1PR1-dependent inhibition of angiogenic sprouting is connected to inhibition of VEGFR2 signaling (57). The authors suggested that S1PR1 suppresses the angiogenic responsiveness in vessels at a developmental stage. Our data are in agreement of their demonstration because we showed the expression of S1PR1 at the adult stage in the lungs when it is important for vessel stabilization, an effect shown to restrict sprouting in at least in part by stabilizing VE-cadherin.β-catenin which is a major partner of VE-cadherin, is well known to activate complexes with transcription factors. We found that FOXF1 is the transcription factor that plays a role in our processes. Among the diverse roles for FOXF1, it is in development that it has been well described since it controls cell cycle progression, cell survival, expression of differentiated genes, and cell metabolism (75). The binding of FOXF1 to *S1PR1* promoter is increased in ECs from KI and this is the first time it is described in relation to VE-cadherin mutation. Moreover, the VE-cadherin is maintained at the membrane which reinforce the junctions.

## SIGNALLING PATHWAYS

In attempt to characterize the potential signalling pathways involved in that process, we search for the major known to that context which is the VEGF-VEGF-R2. Our transcriptomic analysis showed a downregulation of VEGFR2 (VEGFR2 −1.77, p-value=0.00845) which was confirmed at the RNA level and the protein level and was in agreement with the overexpression of S1PR1. The cross talk between a seven-transmembrane-domain G-protein coupled receptor (S1PR1) and a growth factor receptor with tyrosine kinase activity (VEGFR2) might need further studies to determine if such pathways are involved in some of the abnormalities seen in the tumor blood vasculature (77). To date five major autophosphorylation sites within VEGFR-2 have been studied in the past years. Y^951^ lies in the kinase insert domain, Y^1054^ and Y^1059^ are in the kinase domain, whereas Y^1175^ (Y^1173^ in mouse) and Y^1214^ are in the C-terminal portion of the receptor (76). As a result, we found that VEGFR2 protein was decreased in KI mice as well as the phosphorylation on pY^1173^, which strongly suggest a lack in binding partners, and thus an impaired migration process (76). Meanwhile, its ligand VEGF was unchanged thus any effect of VEGF can be partially downregulated because of the dowregulation of VEGFR2 expression in ECs.

Although it was demonstrated that in a quiescent endothelium, no phosphorylation of adherens junctions was found (67,68), in the lung endothelium we found that VE-cadherin was significantly phosphorylated at site Y^731^ more predominantly in KI ECs. These results are in agreement with our previous results in mice (11). The pulmonary capillary is only separated from the alveolar space by a layer of extracellular matrix and an epithelial cytoplasm each around 0.1 mm thick (69). Particularly high VEGF levels are expressed in the lungs, reflecting the critical role of VEGF for lung structural maintenance of the adult lung (70). This might explain the level of phosphorylation of VE-cadherin. The tyrosine kinase c-Src has been shown to directly phosphorylate adherens junction targets, including VE-cadherin, and facilitate adherens junction disassembly. When activated Src can lead to the phosphorylation of Y^658^ and Y^731^-VE-cadherin and thus may account for the well known role that Src kinases play in VE-cadherin-mediated cell-cell junctional activity. In VE-cadherin mutant, we found that VE-cadherin is still phosphorylated at site Y^731^ which disrupt its ability to bind β-catenin (61). As the Y^731^ site is unique to VE-cadherin when compared to other cadherins, it suggests that the regulation of this residue may be involved in an endothelium specific mechanism. Knowing that β-catenin acts as a transcriptional activator, the KI mice could represent an interesting model to understand how β-catenin/VE-cadherin interaction regulates transcription of target genes in specific disease like atherosclerosis, or cancer.

## LUNG PATHO-PHYSIOLOGY IS CONTROLLED BY VE-CADHERIN TYROSINE PHOSPHORYLATION

Our study was performed on ECs from lung and thus we kept in mind to determine if there was any potential effect of lung vessels. We found a reduction in vascular density with larger and thicker vessels, and an decrease in collagen deposit in the lung of KI mice. Moreover, the endothelium must have perform some secretory functions as we could see three proteins in the BAL that were increased in the KI mice. Dll4 is known as a negative regulator of angiogenic sprouting (62), Osteopontin is a calcium-binding glycol-phosphoprotein which is involved in tissue injury and repair (78) and IGFB2 (insulin-like growth factor-binding protein) is related to angiogenesis in tumor context (79). The lung fibrosis are not completely understood and moreover, very few studies examine the role of endothelial cells. Future studies are required to fully understand the molecular mechanism of endothelial dysfunction involved in the fibrosis.

## CONCLUDING REMARKS

We demonstrated that the genetic factors related to the phosphorylation of VE-cadherin are as important as those related to the presence or absence of VE-cadherin but with a particular specificity which led us to the processes of cell adhesion. and angiogenesis including the VEGFR2 pathway (80). We only focused on one gene that we thought was important in the VEGFR pathway which was known to stabilize the vascular endothelium. This is the S1PR1 gene. In the KI mouse, the absence of phosphorylation on the Y^685^ site of VE-cadherin allows the phosphorylation of the Y^731^ site and the delocalization of B-catenin in the nucleus which activates FOXF1 which binds to the promoter from S1PR1. In the case of tumors presenting abnormalities of vessels, it can be considered that the epigenetic due to the phosphorylation of the VE-cadherin leads to a reprogramming of the ECs and this may open up new therapeutic avenues to fight against abnormalities of tumor vessels, to improve blood flow and to increase the accessibility of chemotherapy.

## ACKNOWLEDGMENTS

The authors are indebted to Soumalamya Bama-Toupet, Charlène Magallon and Hervé Pointu for their technical assistance in the animal facility. We also want to thank Aude Salomon and Violaine Simon for their excellent technical assistance. Véronique Colin-Faure was the IRIG flow cytometry plaform manager and we thank her for excellent technical support for cell sorting. Frederic Sergent and Nicolas Lemaitre for their technical expertise regarding immunohistochemistry of lungs. Frederic Mitler and Frederic Kemarec for immunoblot and pcr advices, Patricia Obeid for images quantification. We thank Dr Saporano sfor excellent technical advices for CHIP assays

